# Advancing Codon Language Modeling with Synonymous Codon Constrained Masking

**DOI:** 10.1101/2025.08.19.671089

**Authors:** James Heuschkel, Laura Kingsley, Noah Pefaur, Andrew Nixon, Steven Cramer

## Abstract

Codon language models offer a promising framework for modeling protein-coding DNA sequences, yet current approaches often conflate codon usage with amino acid semantics, limiting their ability to capture DNA-level biology. We introduce SynCodonLM, a codon language model that enforces a biologically grounded constraint: masked codons are only predicted from synonymous options, guided by the known protein sequence. This design disentangles codon-level from protein-level semantics, enabling the model to learn nucleotide-specific patterns. The constraint is implemented by masking non-synonymous codons from the prediction space prior to softmax. Unlike existing models, which cluster codons by amino acid identity, SynCodonLM clusters by nucleotide properties, revealing structure aligned with DNA-level biology. Furthermore, SynCodonLM outperforms existing models on 6 of 7 benchmarks sensitive to DNA-level features, including mRNA and protein expression. Our approach advances domain-specific representation learning and opens avenues for sequence design in synthetic biology, as well as deeper insights into diverse bioprocesses.

## Introduction

Codons are triplets of nucleotides that encode amino acids during the translation of mRNA into proteins. Although there are only 20 standard amino acids, the genetic code comprises 64 codons. This redundancy means that most amino acids are encoded by multiple synonymous codons. As a result, a single protein sequence can be represented by a vast number of different DNA sequences.

Codon optimization is the process of selecting a DNA sequence that maximizes desirable traits, such as expression, without altering the amino acid sequence. Traditional methods often rely on heuristics, such as using the most frequent codons in the host organism or matching codon usage frequencies^1–11^. However, these approaches are limited in scope and do not capture the complex biological factors that influence translation efficiency, folding, and other aspects of gene expression.

Recent work has introduced codon language models to learn patterns in coding DNA using deep learning^12–17^. These models are inspired by protein language models, which have shown remarkable success in structure prediction and mutation modeling^18^. However, current codon language models suffer from a key limitation in how they are trained. During masked language model training, the model is asked to predict a masked codon from the entire codon vocabulary, including codons that encode different amino acids. This means that when a codon for alanine is masked, the model is penalized not only for predicting incorrect synonymous codons, but also for predicting codons that encode completely different amino acid, codons that are biologically invalid in that context. These predictions introduce noise into the learning signal and hinder the model’s ability to focus on codon-level variation.

This training setup introduces a confounding factor. The model must first learn which amino acid belongs at a given position and then which codon is most appropriate for that amino acid. As a result, it becomes difficult to isolate patterns that are specific to codon usage and DNA-level features.

In parallel, techniques such as logit masking and targeted logit manipulation, where certain outputs are deliberately excluded or down weighted during prediction, have emerged as active research areas in the field of machine learning. These methods are particularly useful for enforcing known structural constraints and improving model interpretability^19–22^, in this case, codon optimization where the protein sequence is known *a priori,* and the objective is to optimize the corresponding DNA sequence. Logit masking has been used in generative codon language models to make sure codon output translates to the input protein sequence^23,24^.

However, these models differ fundamentally in their design: one operates on amino acid sequences and does not represent codons directly, while the other applies synonymous constraints only during inference, not during training. Our work extends these ideas to the training objective itself, demonstrating how biologically grounded logit constraints can be integrated into masked language modeling to guide representation learning. This approach aligns with broader efforts in ML to incorporate structured priors and task-specific constraints into foundational models and illustrates how training-time logit-space control can reshape learned representations in biologically meaningful ways.

Building on these insights, we introduce a new training strategy, synonymous codon-constrained masking. When a codon is masked, the model is restricted to predicting only codons that encode the same amino acid. For example, if the masked codon is ‘AGT’, which encodes serine, the model is only allowed to predict serine-encoding codons, ‘AGT’ or ‘AGC’. This constraint is enforced by applying the softmax operation only over the subset of synonymous codons, rather than the full codon vocabulary. As a result, the model no longer needs to infer amino acid identity during training and can focus entirely on learning codon-level semantics.

Our model, trained with this strategy, outperforms existing codon language models on tasks that are sensitive to DNA sequence variation. Furthermore, analysis of the learned embedding space reveals that our model clusters codons based on nucleotide composition, while unconstrained models tend to group codons by amino acid properties, highlighting the conflation of codon usage with protein-level semantics in previous approaches. This makes our model particularly well-suited for applications in nucleotide-based therapeutics, such as mRNA vaccines, gene therapies, and oligonucleotide design. It also provides a powerful tool for codon optimization in biotherapeutics, where synonymous codon choices can significantly affect protein yield, folding, and other sequence-dependent properties, without altering the amino acid sequence. Beyond its biological utility, SynCodonLM illustrates how architectural and objective-level choices can disentangle latent factors in structured sequence modeling, offering insights applicable to other domains with symbolic or hierarchical vocabularies.

## Results

### Motivation for Synonymous Codon-Constrained Training

Large language models (LLMs) developed for proteins have significantly influenced the design of codon language models. While standard masked language modeling (MLM) approach could be used, predicting masked tokens from the full codon vocabulary, we hypothesized that allowing non-synonymous codons in the prediction space introduces a confounding factor.

This issue is exemplified in the CaLM model^13^, which maintained high performance on protein property prediction even when the coding DNA sequence was entirely mutated. This suggests that the model’s performance was driven in some part by amino acid-level information in addition to codon-level features. Such findings reinforce our motivation to constrain the model’s output space to only synonymous codons during training, thereby encouraging the model to learn codon usage patterns independently of protein-level semantics.

### SynCodonLM Architecture and Species-Aware Training

Our model was trained on the largest dataset used to date for any codon language model, comprising over 66 million coding sequences (CDS) sourced from NCBI RefSeq^25^. These sequences span more than 30,000 unique species, offering unprecedented diversity and coverage across the tree of life. As shown in **Figure 1a**, the dataset was grouped into broad organismal categories defined by NCBI taxonomy, with the number of sequences per group visualized to highlight the distribution of training data.

**Fig. 1:**
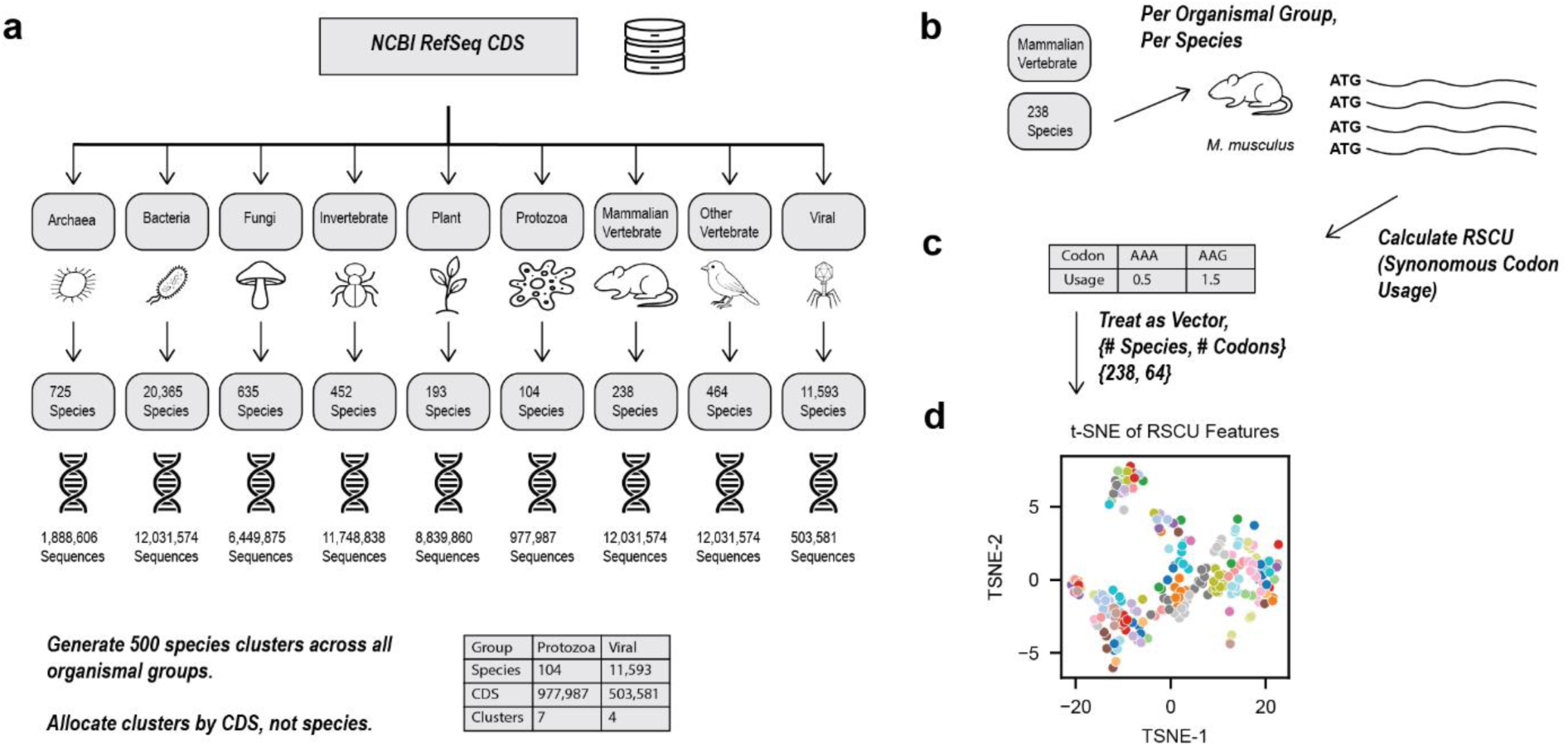
Dataset Curation and Species Clustering **a** Using NCBI RefSeq database, we cycled through all 9 of its major organismal groupings, along with all species (while excluding cell lines and sub-species), to gather codon sequences (CDS). **b** CDS were grouped per organismal group and **c** RSCU was calculated per species. RSCU was treated as a vector across all species in an organismal group and k-means clustering was used to cluster species with similar codon usage. These clusters are visualized in **d** using t-SNE, where species with similar codon usage patterns (colored by cluster) appear closer together in the embedding space.

To capture codon usage patterns across species, we computed relative synonymous codon usage (RSCU)^6^ values for each species within its respective organismal group (**Fig. 1b**). These RSCU vectors were treated as high-dimensional representations of codon bias (**Fig. 1c**) and clustered using k-means. The resulting clusters were visualized using t-SNE to reveal global structure in codon usage across taxa (**Fig. 1d**).

Inspired by the CodonTransformer model^15^, we introduced learnable species-level differentiation into SynCodonLM by incorporating token type embeddings. Specifically, each codon token was embedded by summing a content (codon) embedding with a token type embedding derived from the species’ RSCU cluster (**Fig. 2b**). This approach provides the model with biologically relevant context about codon usage patterns, without requiring species-specific tokens that would dramatically increase the size of the token type embedding matrix. This design balances biological specificity with model scalability, enabling the incorporation of species-level context without inflating the number of learned parameters. The resulting token type embeddings exhibit clustering by biological category of species (Supplementary Figure 1), suggesting that the model captures taxon-specific codon usage patterns.

**Fig. 2:**
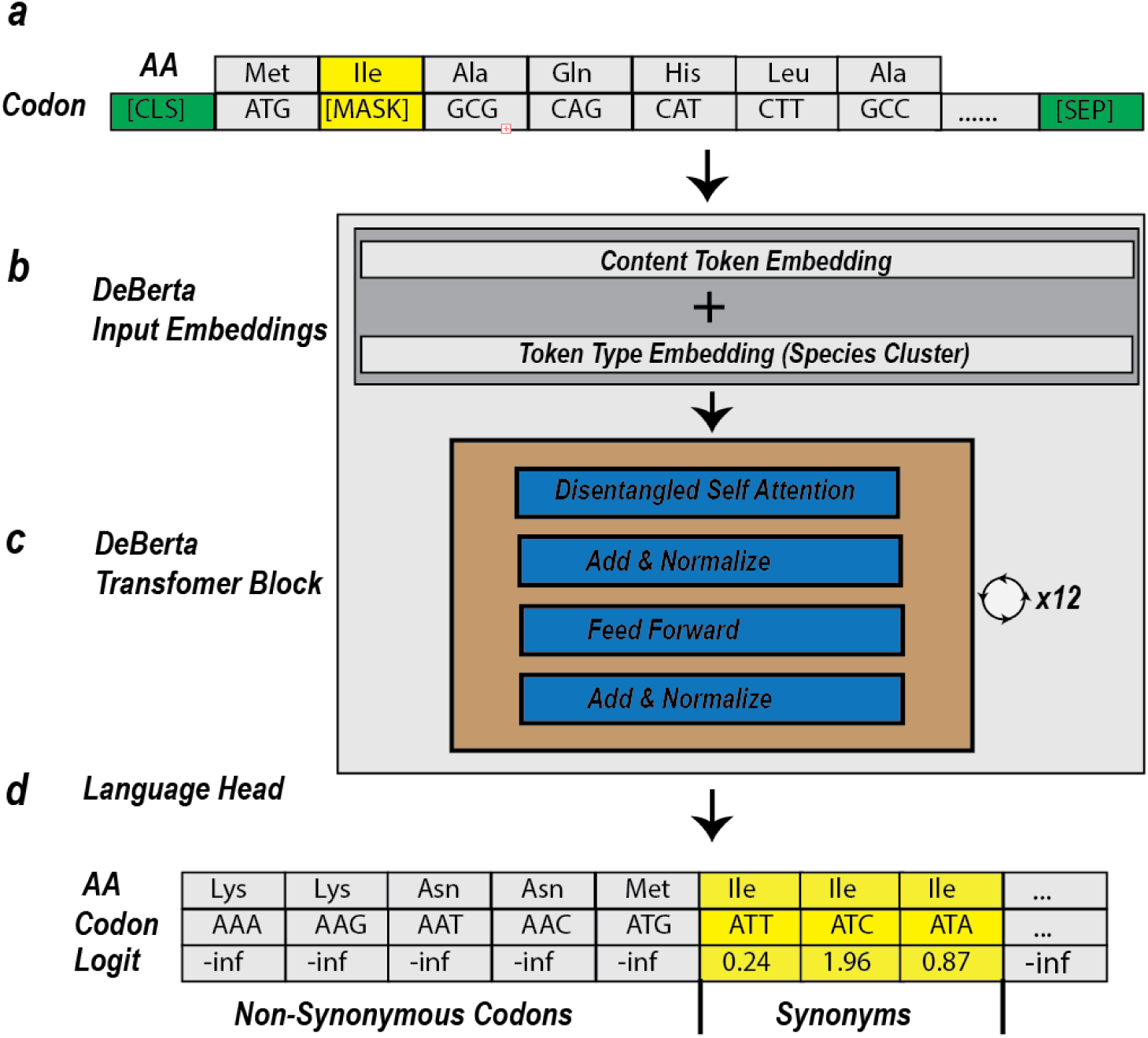
Constrained Decoding of Model Output During Training. **a** SynCodonLM masks roughly 15% of input codons. **b** Token inputs receive their static, respective embeddings, which have a token type embedding added based on the species corresponding species group. **c** Input embeddings are fed into the DeBerta transformer block, where they have disentangled self-attention applied utilizing relative positional embeddings. Our model has 12 layers of this transformer block. **d** Model output logits are masked with -inf for positions which correspond with non-synonymous codons.

During training, 15% of codons were randomly masked (**Fig. 2a**), and the model was tasked with predicting the masked codon using a DeBERTa-style transformer architecture (**Fig. 2c**). DeBERTa (Decoding-enhanced BERT with disentangled attention) represents a state-of-the-art advancement in BERT-based models, achieving top performance across a wide range of natural language understanding (NLU) benchmarks^26^. Its disentangled attention mechanism and enhanced position encoding allow for more expressive and efficient representation learning, making it particularly well-suited for capturing subtle patterns in biological sequences. Training convergence was stable across epochs, as shown in Supplementary Figure 2.

Crucially, we applied a synonym-aware logit masking strategy at the output layer (**Fig. 2d**), where logits corresponding to non-synonymous codons were set to negative infinity. This biologically grounded constraint ensures that the model learns codon-level variation independent of amino acid identity. The effectiveness of this constraint is reflected in the MLM accuracy, which exceeds 60% during training (Supplementary Figure 3).

### SynCodonLM Captures DNA-Level Features, Not Protein Semantics

To validate that our synonym-constrained training strategy produces a model that focuses on coding DNA characteristics rather than protein-level semantics, we analyzed the learned input embeddings of all codon tokens in the vocabulary.

Previous codon language models^12–16^ have shown that codon embeddings tend to cluster based on amino acid properties. In Supplementary Figure 4a-e, we performed a t-SNE on the input embeddings of 5 other codon language models, which consistently reveal clustering by amino acid identity or physiochemical property. These models were trained without synonymous constraints, and their clustering by amino acid identity indicates an entanglement of codon and protein semantics in the learned representations. This suggests a confounding influence of protein-level information, likely arising from unconstrained training objectives that require the model to infer the correct amino acid before selecting a codon. In contrast, our model, SynCodonLM, was trained to focus exclusively on synonymous codon variation, allowing it to learn patterns intrinsic to DNA.

When visualizing the input embeddings of SynCodonLM using t-SNE, we observed that codons clustered based on nucleotide-level features. In **Figure 3a**, codons are grouped by their wobble base, which is the third nucleotide in the codon. This clustering reflects biologically relevant structure, as the wobble position plays a critical role in tRNA recognition and codon degeneracy^27^, has been shown to influence translation speed^8,9,28^ and mRNA stability^29^.

**Fig. 3:**
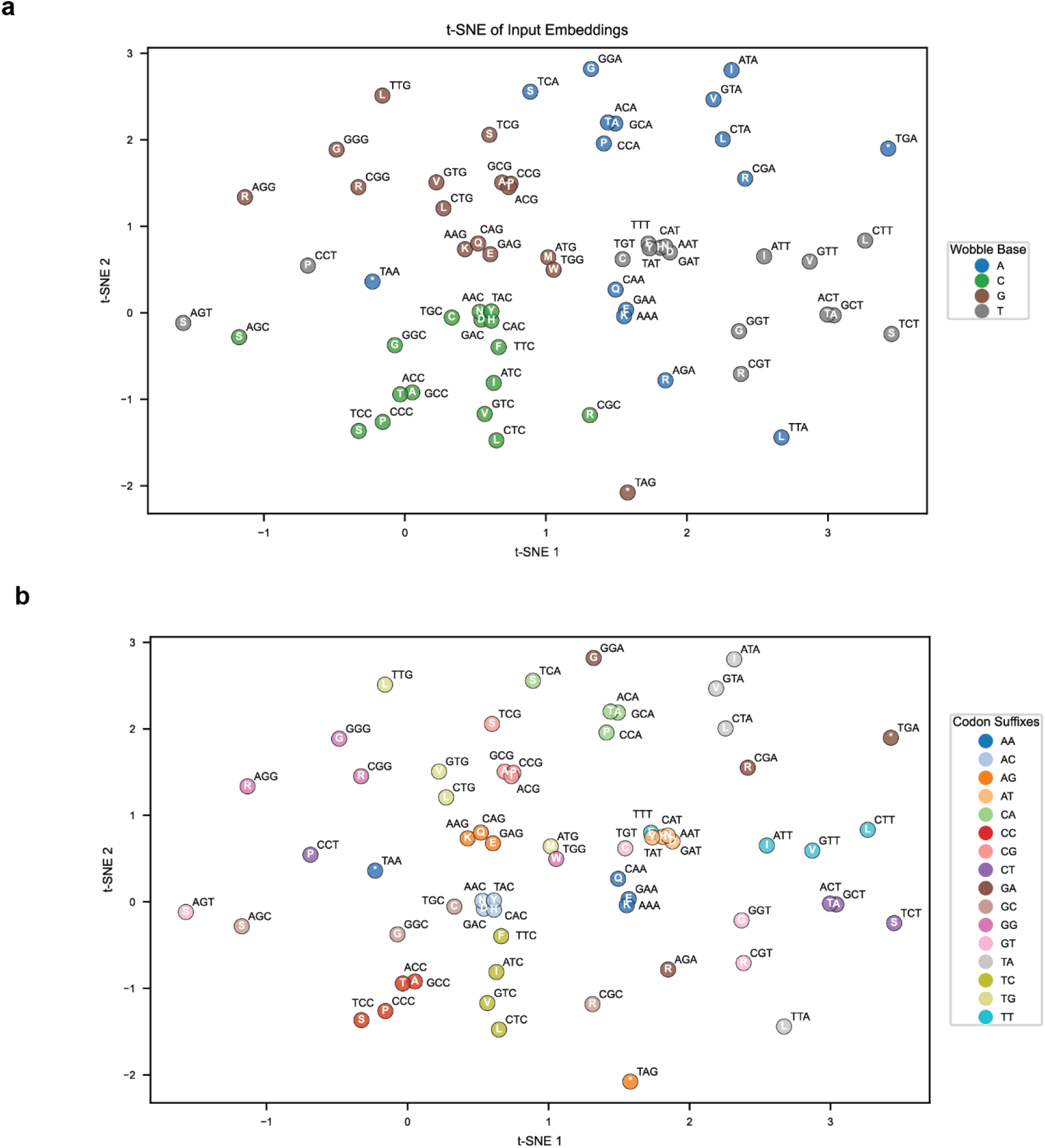
SynCodonLM Learned DNA Specific Features Within Codons **a** t-SNE projection of input embeddings for all codons reveals that codons cluster according to their third (wobble) base. Each point is colored by its wobble nucleotide (A, C, G, or T), highlighting the influence of the wobble position on the learned embedding space. Separation by wobble base is statistically significant (PERMANOVA, pseudo-F=3.025, p < 0.001). **b** When colored by codon suffix dinucleotide (positions 2 and 3), the same t-SNE projection shows that codons with similar suffixes tend to group together, suggesting that the model captures dinucleotide-level patterns in codon representation. Separation by suffix dinucleotide is statistically significant (PERMANOVA, pseudo-F=1.937, p < 0.001).

In **Figure 3b**, codons are grouped by their dinucleotide suffix, which consists of the last two nucleotides of the codon. Codons with the same suffix cluster together regardless of the amino acid they encode. This clustering reveals further biologically relevant structure, as there have been findings that suffix dinucleotides can induce unique structural changes in mRNA^30^, and that these structural features can significantly alter translation dynamics^31,32^.

Both clustering patterns were statistically significant, as confirmed by PERMANOVA tests with a p-value less than 0.001. Additionally, codons that do not have any synonymous alternatives, such as ‘ATG’ for methionine and ‘TGG’ for tryptophan, appear near the center of the t-SNE plots in **Figures 3a** and **3b**. This suggests that the model did not learn strong embedding features for these codons, which is expected given their lack of synonymous variation. Most importantly, there is no evidence of clustering by amino acid identity, which supports the conclusion that SynCodonLM learns nucleotide-driven representations rather than protein-driven ones.

These embedding patterns are a direct consequence of our logit-constraining strategy during training. By masking logits corresponding to non-synonymous codons, the model is prevented from learning amino acid-level distinctions and instead focuses on nucleotide-level variation. This constraint reshapes the optimization landscape, guiding the model toward codon-specific representations. In contrast, models trained without logit masking tend to encode semantic relationships between codons and their corresponding amino acids, as reflected in their embedding clusters.

### SynCodonLM Embeddings Outperform Existing Models on Codon-Sensitive Tasks

To assess the utility of the embeddings produced by SynCodonLM, we evaluated model performance across multiple datasets in which codon usage varies while the encoded protein sequence remains constant. These datasets were specifically chosen to reflect biologically meaningful differences in mRNA or protein properties that arise from synonymous codon variation. This contrasts with prior studies, where evaluation datasets often include differences in both coding DNA and protein sequence, making it difficult to disentangle codon-level effects from protein-level semantics.

This distinction is critical. Models such as CaLM^13^, cdsBERT^12^, CodonBERT^14^, CodonTransformer^15^ and Mistral Codon^16^ are trained using unconstrained objectives that allow them to learn a mixture of codon and protein language. As a result, their embeddings may reflect amino acid properties rather than DNA-level features. In contrast, SynCodonLM was trained using a synonym-constrained masking strategy, ensuring that its embeddings capture codon-specific signals independent of protein identity.

As shown in **Figure 4a**, we performed five-fold cross-validation on each dataset, repeating the process across 20–80 random seeds. For each seed, mean-pooled embeddings served as input to a linear regression head trained to estimate the fitness metric for each dataset. Average R² scores were then computed across folds and these averages were used to calculate the mean performance per model and dataset, and to perform paired t-tests comparing SynCodonLM against other models.

**Fig. 4:**
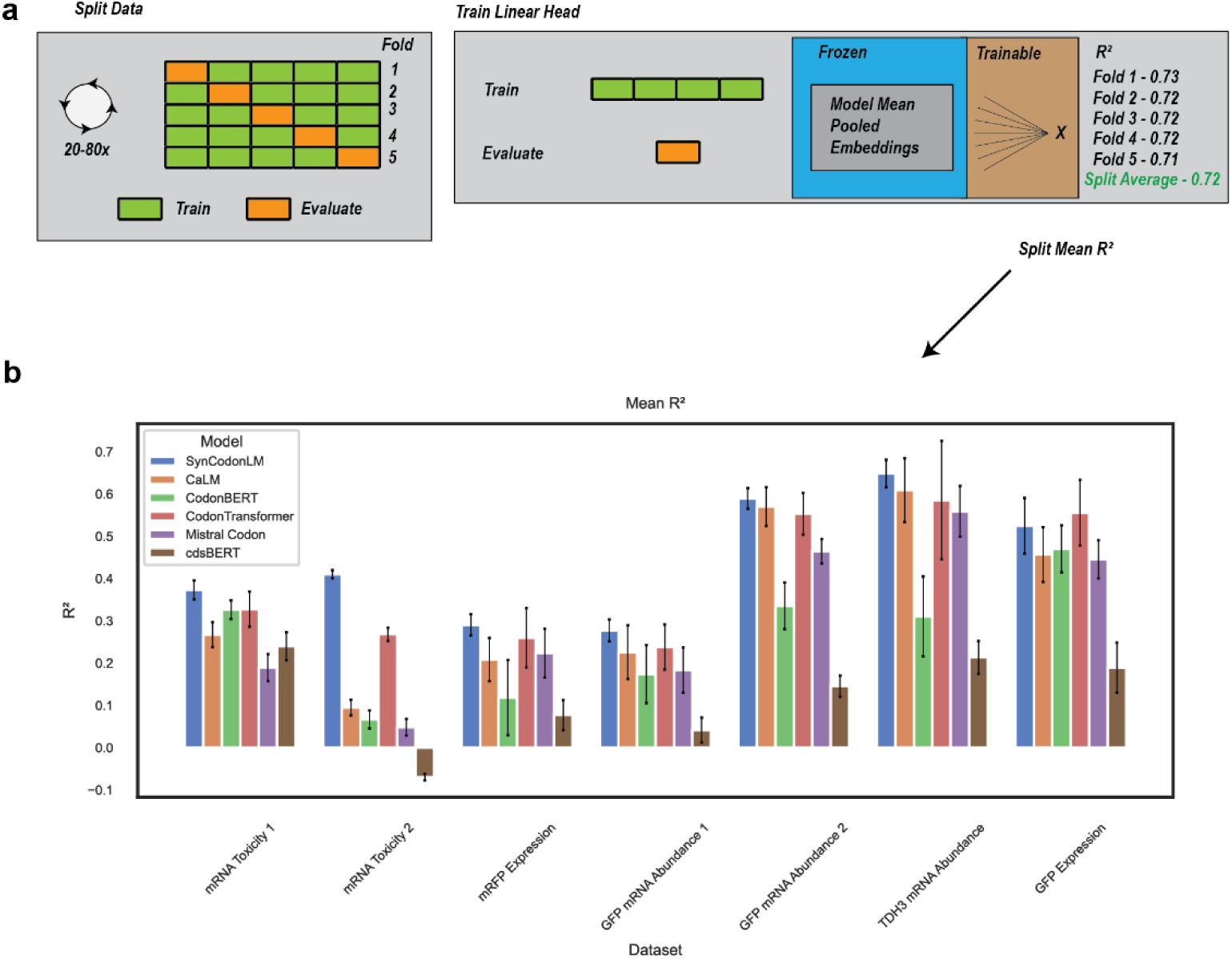
SynCodonLM Outperforms Other Models **a** We performed 5-fold cross validation on each dataset, training a linear head on mean pooled embeddings from each model, for each of the 5 folds. For each set of 5 folds, we aggregated the split’s evaluation metrics. We re-split data 20-80 times to get 20-80 average R^2^ for each split, per model, dataset. **b** SynCodonLM outperforms all 5 other codon language models in 6 out of 7 datasets (*p* < 0.05, Paired-T test). SynCodonLM performs better than 4 out of 5 other codon language models in another dataset (*p* < 0.05, Paired-T test).

The results, shown in **Figure 4b**, demonstrate that SynCodonLM achieved the highest performance in six out of seven datasets, significantly outperforming all five other evaluated codon language models (*p* < 0.05, paired t-test). In the remaining dataset, SynCodonLM ranked second, outperforming four out of five models (*p* < 0.05, paired t-test)..

These findings support the assertion that SynCodonLM learns biologically meaningful DNA-level representations, making it better suited for tasks where synonymous codon variation drives functional outcomes. These results demonstrate that constraint-aware training can yield more interpretable and biologically faithful representations, offering a framework for structured sequence modeling in other domains.

## Discussion

In this work, we introduced SynCodonLM, a codon language model trained with a synonym-constrained masked language modeling strategy. SynCodonLM is the first codon language model designed to learn nucleotide-level patterns in coding DNA, without being confounded by protein-level semantics. This distinction is evident in the learned input embeddings, which cluster codons based on nucleotide features such as wobble base and suffix dinucleotides, unlike prior models that group codons by amino acid identity.

By removing the need to infer amino acid identity during training, SynCodonLM captures codon-level variation that is both interpretable and predictive. It consistently outperforms existing codon language models on tasks where synonymous codon changes drive functional outcomes, particularly when the protein sequence is held constant.

Beyond our novel training objective, SynCodonLM benefits from two key innovations: it leverages the DeBERTa architecture for enhanced representation learning, and it is trained on roughly 60 million coding sequences from nearly 35,000 species, the largest and most diverse dataset used in codon language modeling to date.

Central to our approach is the use of logit-space constraints during training, a technique increasingly recognized in machine learning for enforcing structural priors and improving interpretability. By masking logits for non-synonymous codons, we align the model’s prediction space with biological reality, guiding it toward disentangled representations. This strategy reflects a broader principle in ML: that domain-specific constraints can reshape foundational model training to better capture latent structure. Similar approaches are emerging in symbolic reasoning, chemical modeling, and grammar-constrained generation, where output validity and semantic disentanglement are critical.

SynCodonLM has broad potential for downstream applications, especially in nucleotide-based therapeutics and biotherapeutics. In these domains, silent mutations are often used to alter biological properties. For example, codon optimization has been shown to increase vaccine efficacy^33–35^, enhance viral fitness in oncolytic adenoviruses^11^, increase mRNA stability and translation efficiency in AAV payloads^36,37^, improve AAV capsid packaging^38^, boost therapeutic antibody yield^39–41^ and influence post-translation modifications^42^.

SynCodonLM could also be applied to large-scale public datasets, such as those from the Protein Abundance Database^43^, to build predictive models for codon optimization. Its ability to isolate the contribution of coding sequence, independent of protein-level signals, could enable species-specific models that reveal new insights into translational regulation.

Together, these capabilities position SynCodonLM not only as a powerful tool for codon-aware modeling, but also as a case study in how constraint-aware training can enhance representation learning across structured domains in biology and beyond.

## Methods

### Pre-Training Data

We curated a comprehensive dataset of coding sequences (CDS) from the NCBI RefSeq database^25^, encompassing nine major organismal groups: Archaea, Bacteria, Fungi, Invertebrate, Plants, Protozoa, Mammalian Vertebrates, Other Vertebrates and Viruses. To ensure broad taxonomic coverage and minimize sampling bias, we included only one representative CDS dataset per species, explicitly excluding sub-species and cell line-specific entries. This strategy helped prevent overrepresentation of well-studied organisms and ensured a more balanced view of codon usage across the tree of life.

To ensure high data quality, we restricted our selection to CDS datasets labeled as “reference”, which are curated and represent the most complete and accurate genomic assemblies available in RefSeq^25^. These reference sequences are manually reviewed and serve as gold-standard annotations for genomic studies.

Given the disproportionate abundance of CDS entries in certain groups (e.g., Bacteria and Other Vertebrate), we applied stratified sampling to balance the dataset, with an intentional emphasis on mammalian vertebrates to support downstream modeling objectives. This emphasis reflects the relevance of mammalian systems in biotherapeutics and mRNA-based therapeutics, where codon optimization plays a critical role. A breakdown of dataset composition per organismal group can be seen in **Figure 1a**.

In total, the final dataset comprised 66,503,469 CDS entries from 34,769 unique species. We randomly partitioned the dataset into 90% training (59,853,123 sequences) and 10% testing (6,650,346 sequences) subsets. This large-scale dataset provides unprecedented diversity for codon modeling and is approximately six times larger than that used in any previously published codon language model^12,13,15,17^. All sequences were validated to ensure they contained only canonical nucleotides (A, C, G, T), were divisible by three and contained no internal stop codons.

### Model Architecture

SynCodonLM was trained as a masked language model, based on the DeBERTaV2^26^ architecture, tailored for learning representations of CDS. We utilized DeBERTaV2 due to its demonstrated performance improvements over traditional BERT models^26^, which have been used in other codon language models^14,17^. We selected DeBERTa for its disentangled attention mechanism and relative positional encoding, both of which are well suited for capturing the complex, context dependent relationships inherent in CDS.

Unlike BERT, which adds absolute positional embeddings directly to token embeddings, DeBERTaV2 separates content and positional information within its attention mechanism. This disentangled attention allows the model to better generalize across varying sequence lengths and capture long range dependencies. Additionally, DeBERTaV2 employs relative positional embeddings, which encode the relative distance between tokens rather than their absolute positions. This is particularly advantageous for CDS, where codon context often depends on local and distal relationships within the sequence, such as regulatory motifs or translational hotspots.

We initialized the model with a hidden size of 768, intermediate size of 3072, 12 hidden layer and 12 attention heads and a max length of 1024 positions. This configuration is consistent with that of other codon language models^13–15^. The model was implemented using HuggingFace Transformers (version 4.48.3) and trained using PyTorch Lightning (version 2.4.0) and PyTorch (version 2.7.1+cu126) with mixed precision to optimize memory usage and training speed.

A custom tokenizer was built to include all 64 codons and 5 special tokens. During tokenization, each CDS is prepended with a [CLS] token and terminated by a [SEP] token to denote sequence end. We incorporated token type embeddings into our input representation to give context relative to species for the model to make more informed decisions. While prior works such as CodonTransformer did this per species, our dataset comprises nearly 35,000 species, therefore, the size of the token type id embedding would become extremely large. Meanwhile, their dataset only trained on roughly 150 species^15^. Because of this, we limited token type IDs to a max of 501. This design balances biological specificity with computational scalability, allowing the model to incorporate species-level codon bias without inflating the token type embedding matrix.

As taxonomic groups in our dataset did not have relative sequence to species ratios, we decided to breakdown the allocation of those 501 token type ID by the number of CDS per taxon. To group species together within each taxon, we calculated the RSCU for each species and then performed k-means clustering to assign token type ID across species with similar relative codon usage. This clustering strategy ensures that species with similar translational mechanisms are embedded similarly, improving the biological relevance of the learned representations. A final token type ID was reserved as ‘unknown’ to allow model usage without species-specific embeddings.

### Model Training

The model was trained using an AdamW optimizer with a learning rate warmup from 0 to 2×10^-4^ over the first 10% of steps, followed by a cosine decay back to 0. A dropout rate of 0.1 and weight decay of 0.01 were applied to reduce overfitting and improve generalization. These hyperparameters were selected based on prior work in transformer-based biological sequence modeling.

To increase biological specificity to CDS, we constructed a custom masking matrix of shape [*V*,*V*] where *V* is the vocabulary size, mapping each tokenized codon to only synonymous codons. During training, for each masked token, the corresponding row from the matrix is added to the models’ logits before softmax and computing the loss. This ensured that the model could only predict codons encoding the same amino acid as the masked token.

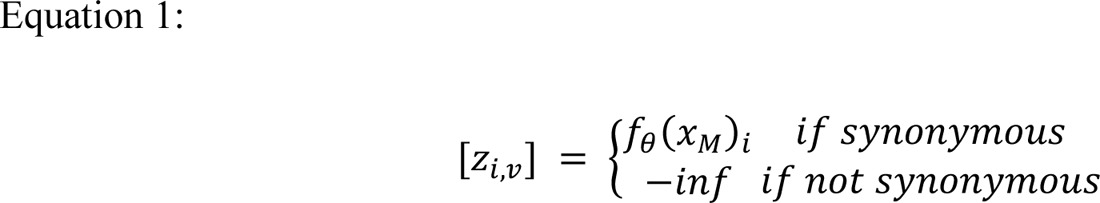

Here, [*Z*_*i,v*_] is the raw logits at position *i* at vocab dimension *v*. The models’ outputs at this masked position *f*_θ_(*x*_*M*_)_*i*_ is retained only for synonymous codons, while logits for non-synonymous codons are set to negative infinity, effectively removing them from the prediction space.

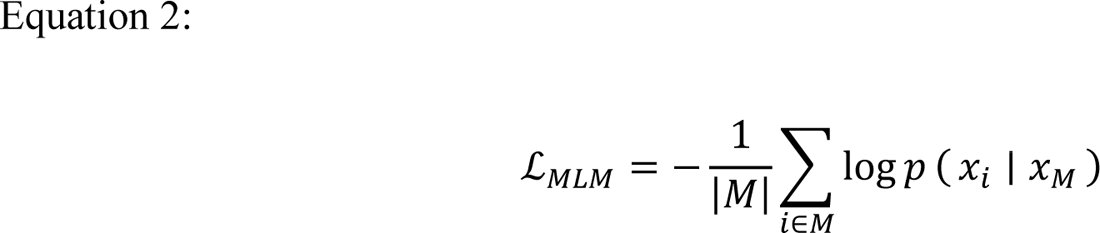

In this formulation, ℒ_*MLM*_ is the masked language modeling loss, computed as the average negative log likelihood over a set of masked positions *M*. For each masked position *i* ∈ *M*, the model predicts the original token *x*_*i*_ given the masked input *x*_*M*_. The loss is normalized by the number of masked positions |*M*| to account for variability in sequence length and masking density.

This synonym-constrained training strategy restricts the model’s language space to biologically valid codon substitutions and prevents penalization for predicting codons that encode different amino acids. By aligning the training objective with biological constraints, the model learns codon-level variation independent of protein semantics. Accuracy was monitored over masked positions throughout training to ensure convergence.

The model was trained with an effective batch size of 1,560, distributed across 12 NVIDIA RTX A6000 GPUs. We utilized 16-bit mixed precision training in PyTorch Lightning to reduce memory usage and accelerate computation. Training was conducted over approximately 12 days, spanning 2 epochs and 76,734 total steps. With an average of 473 codons per sequence, the model processed approximately 56.6 billion tokens, making it one of the largest codon language model training efforts to date.

### Input Embedding Evaluation

We evaluated the static input embeddings learned by SynCodonLM by passing all 64 codon tokens through the model and extracting their corresponding embeddings from the input layer. These embeddings reflect the model’s initial representation of codons prior to any contextual transformation and are useful for assessing how codon-level features are encoded.

To visualize the structure of the embedding space, we applied t-distributed stochastic neighbor embedding (t-SNE) using sci-kit learn^44^ (version 1.5.1) and numpy^45^ (version 1.26.4) to reduce the 768-dimensional embeddings to two dimensions. t-SNE is a non-linear dimensionality reduction technique that preserves local structure and is well-suited for visualizing high-dimensional biological data. We used default t-SNE settings for all visualizations.

The same procedure was applied to generate Supplementary Figure 4a-e which shows other codon language models’ embedding space. This comparative analysis highlights differences in how models encode codon relationships, particularly with respect to nucleotide-level versus amino acid-level clustering.

Additionally, we performed t-SNE on the learned token type embeddings corresponding to species clusters. These embeddings were derived from the RSCU-based k-means clustering described earlier and reflect species-level codon usage patterns. Visualizing these embeddings allowed us to assess whether the model captured biologically meaningful relationships between species based on codon bias.

### Evaluation Datasets

We curated a set of evaluation datasets in which different coding DNA sequences (CDS) encode the same protein but result in measurable differences in biological properties. These datasets were specifically chosen to assess the model’s ability to capture codon-level variation independent of protein sequence, aligning with SynCodonLM’s design objective. By holding the amino acid sequence constant, these tasks isolate the functional impact of synonymous codon changes.

First, we included two datasets from a study that observed mRNA toxicity in *E. coli* when codon-optimizing the same GFP protein^46^. Another dataset demonstrated significant variability in GFP expression in *E. coli* resulting from changes to the first eight codons of the sequence^47^. A fourth dataset reported differential mRFP expression in *E. coli* following random synonymous codon substitutions throughout the sequence^48^.

We also incorporated three datasets from *S. cerevisiae* that measured mRNA abundance after codon optimization of two regions of GFP and one region of the endogenous gene TDH3^10^. These datasets span both prokaryotic and eukaryotic systems and reflect a range of biological outcomes, including expression level, toxicity, and mRNA stability.

Evaluation dataset attributes are summarized in **Supplementary Table 1**. All datasets were min-max scaled to preserve their original variance and comparability across models, except for one dataset with extreme skew toward a central mean and distinct outliers, which was scaled using the Yeo-Johnson transformation implemented in scikit-learn^44^ (version 1.5.1). This preprocessing ensured that model comparisons were not biased by differences in scale or distribution.

### Fine Tuning and Benchmarking

To benchmark SynCodonLM against existing codon language models, we performed fine-tuning on our evaluation datasets using five widely adopted implementations: CaLM^13^, CodonBERT^14^, CodonTransformer^15^, cdsBERT^12^ and Mistral-Codon^16^. We also attempted to include EnCodon^49^, but were unable to extract embeddings due to unresolved bugs in its codebase, which were documented in its GitHub issue tracker.

All models had an embedding dimension of 768, except for cdsBERT, which uses a 1024-dimensional embedding space. To ensure comparability across models, we projected cdsBERT’s embeddings to 768 dimensions using Sparse Random Projection (SRP)^50^, implemented in scikit-learn (version 1.7.1). SRP was chosen over principal component analysis (PCA) because several evaluation datasets contained fewer samples than dimensions, violating PCA’s requirements. This dimensionality reduction allowed for consistent downstream processing across all models.

Given the relatively small size of the evaluation datasets, we froze all base layers of each model and applied mean pooling across the sequence dimension to obtain a single embedding per CDS. A linear regression head was trained on these pooled embeddings using mean squared error loss, with both L1 and L2 regularization to prevent overfitting. We used a batch size of 16 and trained for 100 epochs per model-dataset pair.

To ensure robust performance estimates, we performed five-fold cross-validation and repeated the process across 20 to 80 random seeds, depending on dataset size. This repetition enabled the generation of paired samples for statistical testing, allowing us to compare SynCodonLM’s performance against other models using paired t-tests. Mean performance metrics across folds and seeds were reported for each model and dataset.

### AI Usage

We utilized Microsoft Copilot (GPT-4)^51^ to assist with drafting and refining portions of the manuscript text, including figure legends, supplemental descriptions, and select sections of the Methods. The model was used to improve clarity, consistency, and scientific tone, and to streamline the writing process. All content generated with AI assistance was thoroughly reviewed, edited, and approved by the authors to ensure accuracy and integrity.

## Code Availability

The code used for pretraining, as well as future applications of the model, is available on GitHub at https://github.com/Boehringer-Ingelheim/SynCodonLM. The trained model weights are available on Hugging Face at https://huggingface.co/jheuschkel/SynCodonLM.

## Data Availability

The dataset used for pretraining the model is available on Hugging Face at https://huggingface.co/datasets/jheuschkel/cds-dataset. All datasets used for model benchmarking are available on GitHub at https://github.com/Boehringer-Ingelheim/SynCodonLM/benchmarking-datasets.

## Acknowledgements

We thank Boehringer Ingelheim for providing the high-performance computing resources necessary for model training. J.H. gratefully acknowledges funding support from S.C. through his institute professor endowed chair, as well as additional support from Boehringer Ingelheim.

## Supplementary Information

**Supplementary Figure 1:**
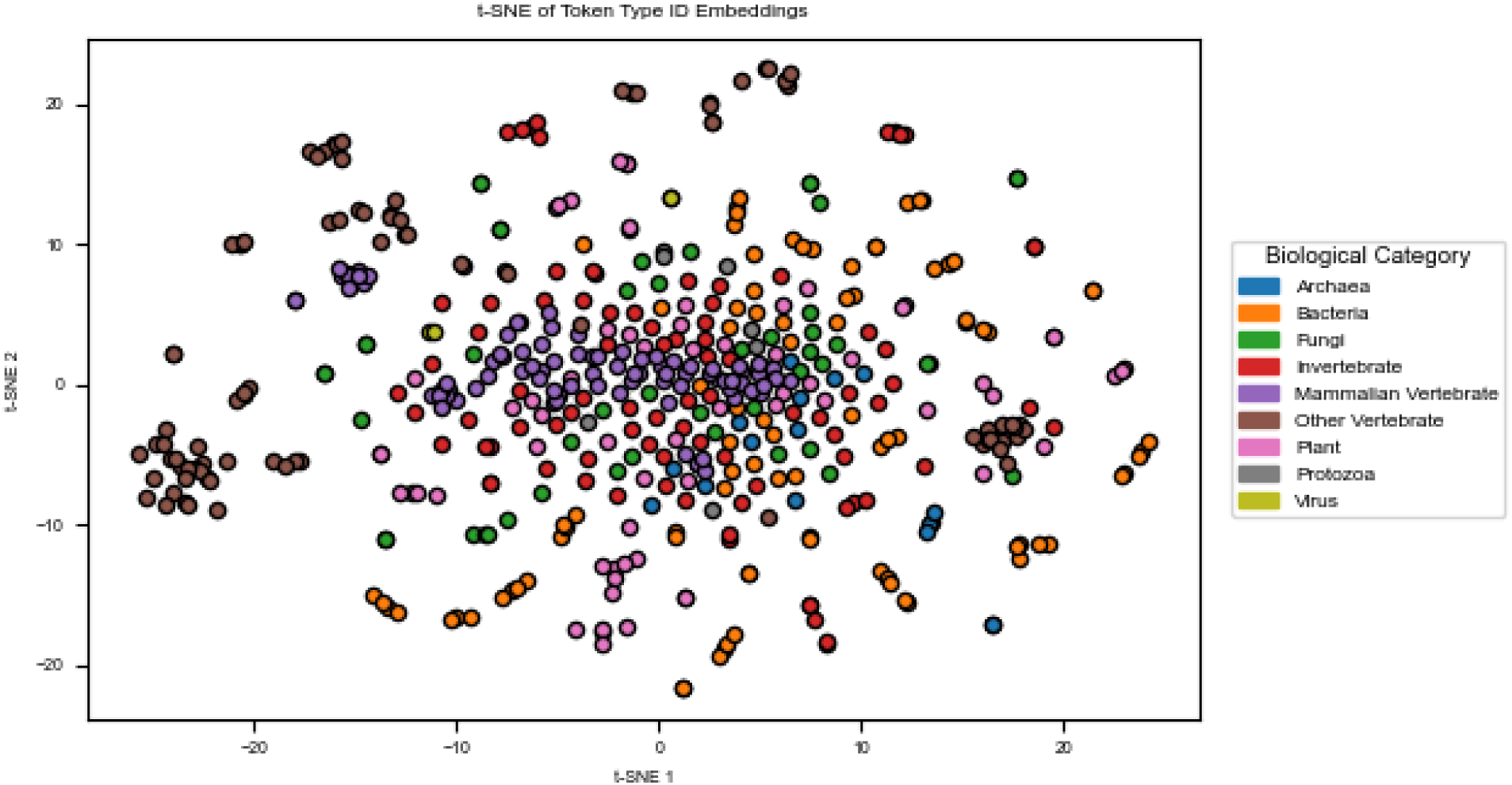
t-SNE of token type ID embeddings shows grouping within biological categories. SynCodonLM appears to have learned some meaningful token type ID input embeddings, as biological categories are visibly grouped near each other.

**Supplementary Figure 2:**
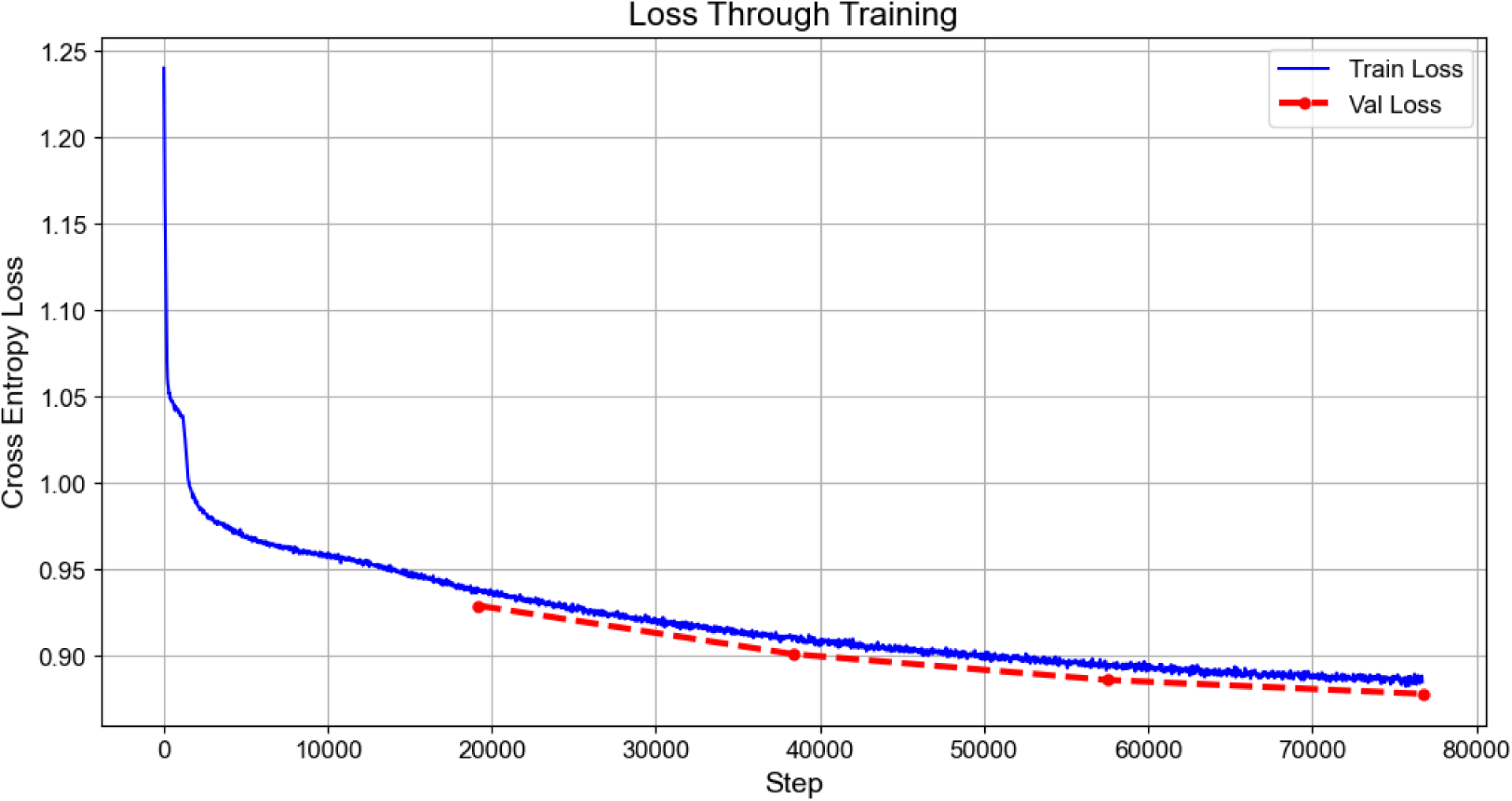
Loss curve through training shows stable convergence.

**Supplementary Figure 3:**
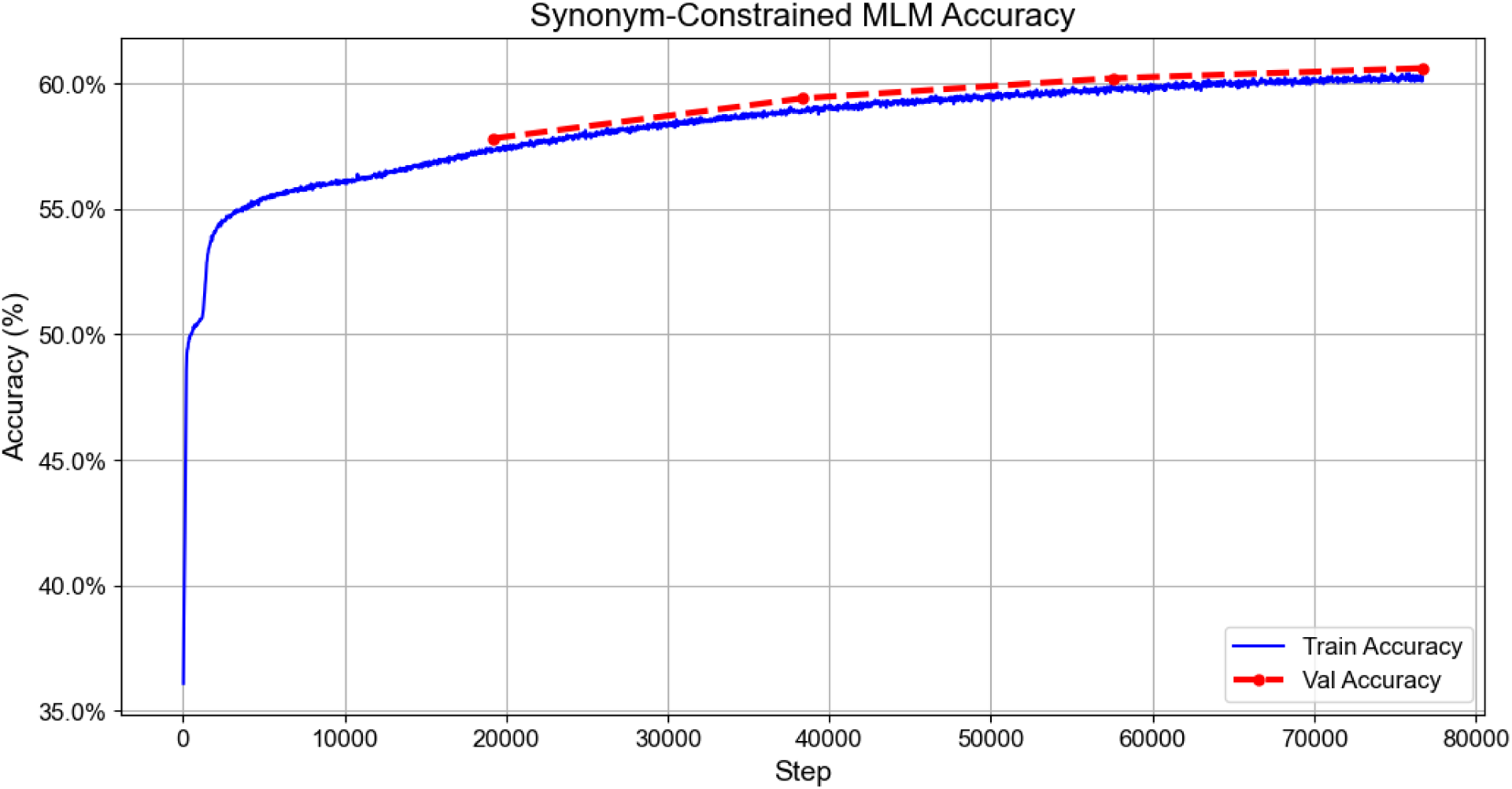
Synonym-constrained masked prediction accuracy through training shows accuracy of over 60% on masked codon prediction.

**Supplementary Figure 4:**
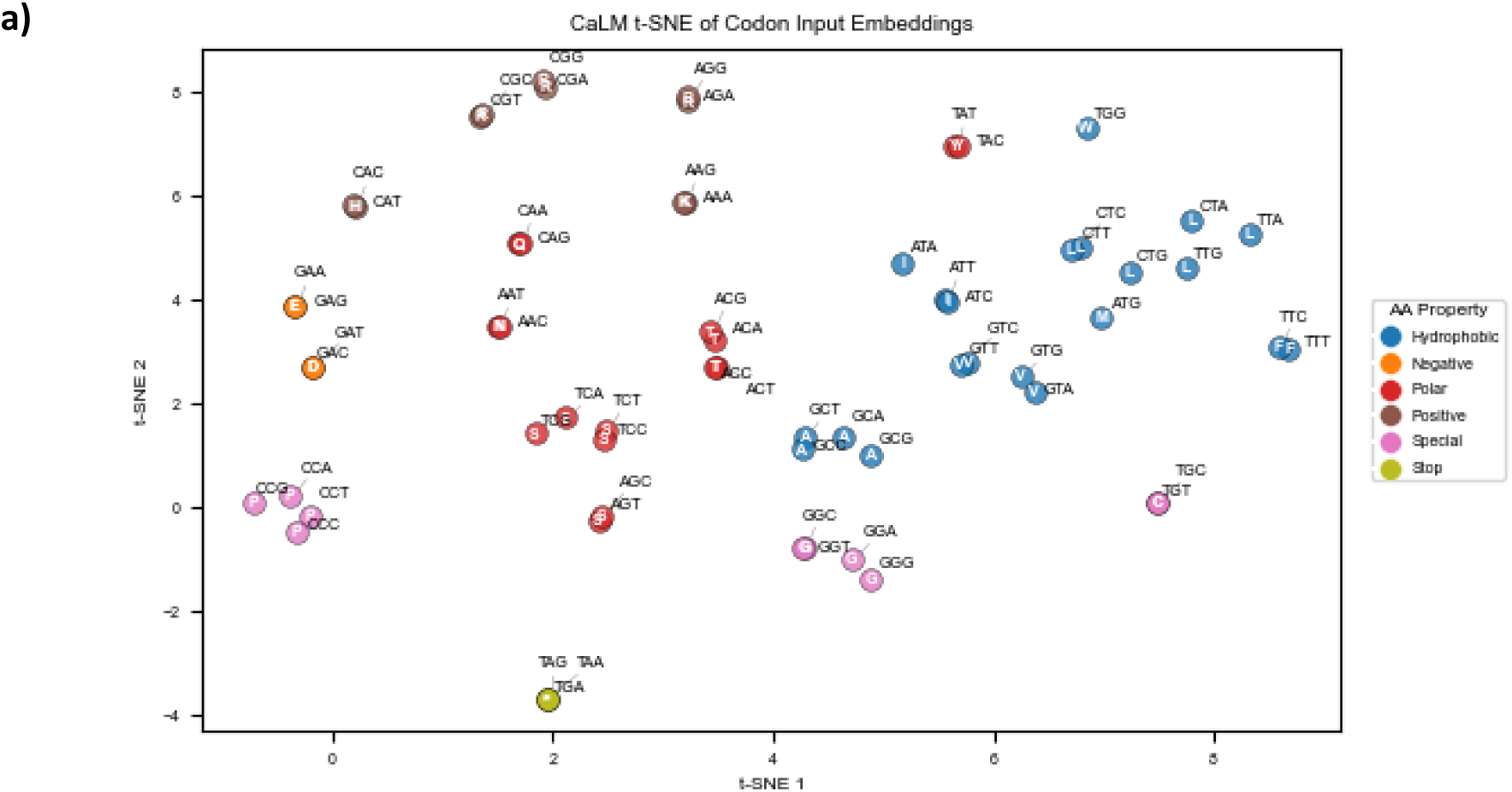
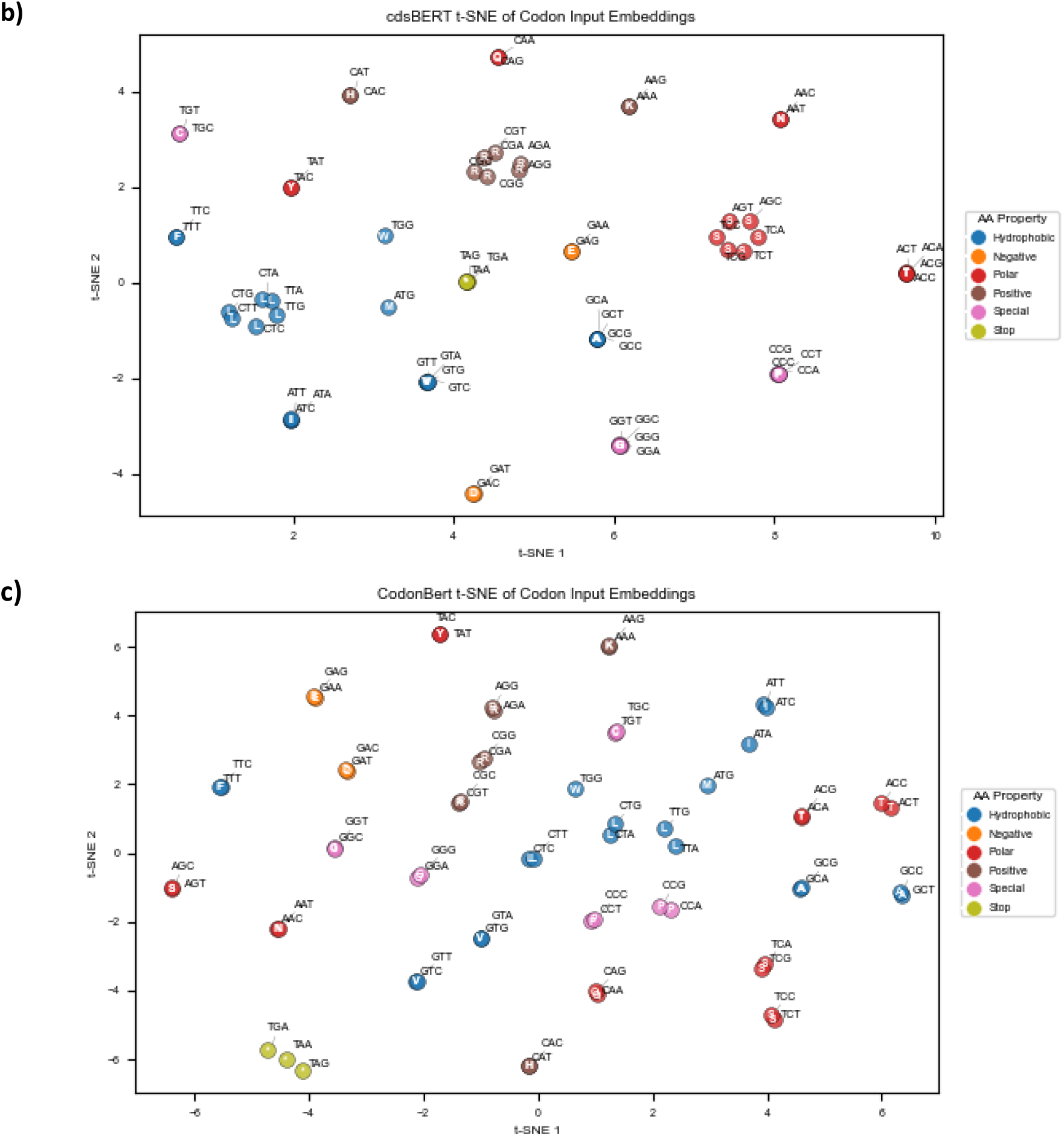
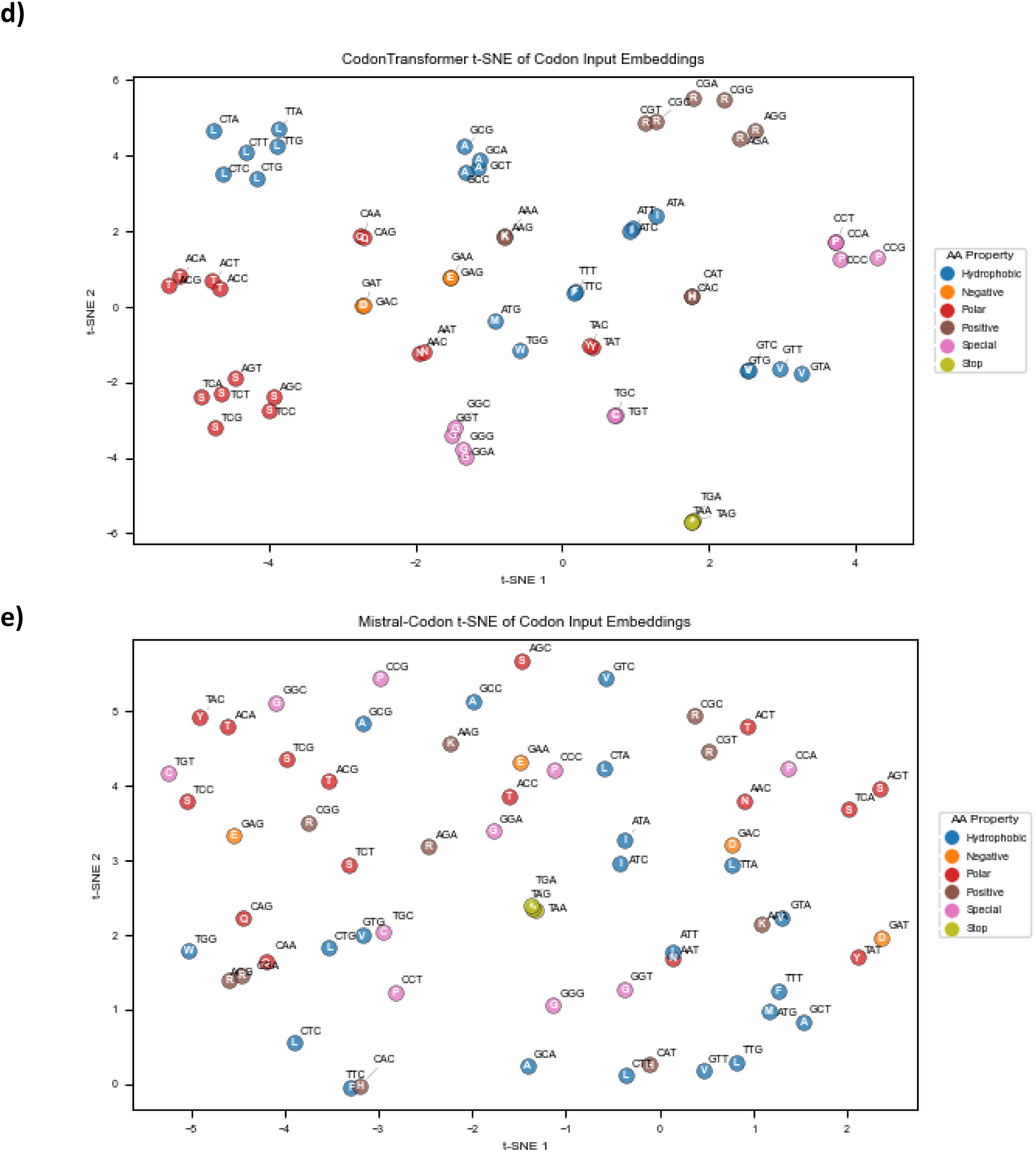
t-SNE of input-codon ID embeddings using (a) CaLM, (b) cdsBERT, (c) CodonBERT, (d) CodonTransformer, and (e) Mistral Codon. In all models, codons are visibly grouped by amino acid/amino acid properties.

**Supplementary Table 1:**
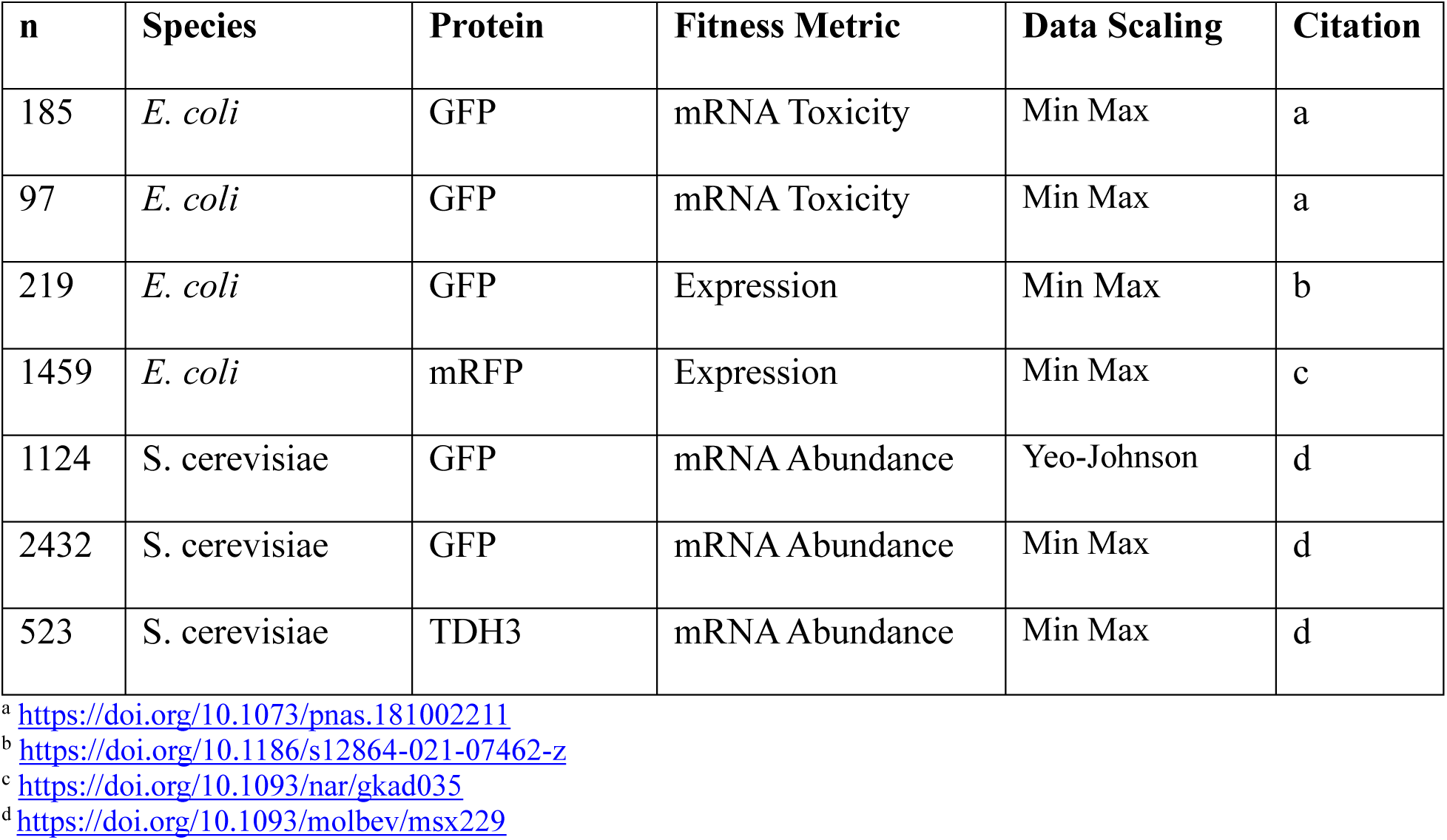
Evaluation dataset properties.

